# Viral mimicry redirects immunosuppressed colorectal tumour landscapes towards a proinflammatory and CMS1-like regenerative state

**DOI:** 10.1101/2024.11.28.625928

**Authors:** Shania M Corry, Svetlana Sakhnevych, Noha Ehssan Mohamed, Sudhir B Malla, Ryan Byrne, Andrew Young, Raheleh Amirkhah, Courtney Bull, Andrea Lees, Keara Redmond, Tamsin Lannagan, Rachel Ridgway, Fiona R Taggart, Natalie C Fisher, Tim Maughan, Mark Lawler, Andrew Campbell, Simon J Leedham, Aideen E Ryan, Dan B Longley, Donna Small, Owen J Sansom, Philip D Dunne

**Affiliations:** The Patrick G Johnston Centre for Cancer Research, Queen’s University Belfast, UK; Cancer Research UK Scotland Institute, Glasgow, UK; University of Liverpool, Liverpool, UK; Wellcome Centre for Human Genetics, Oxford, UK; Discipline of Pharmacology & Therapeutics, School of Medicine, College of Medicine, Nursing and Health Sciences, University of Galway, Ireland

**Keywords:** Immune activation, tumour microenvironment, colorectal cancer, stem cell, therapeutic

## Abstract

In colorectal cancer (CRC), tumours classifier as consensus molecular subtype 4 (CMS4) have the worst prognosis and derive negligible benefit from chemotherapy. We previously described how repressed interferon-related signalling is associated with increased relapse in CMS4 tumours. Although the viral mimetic poly(I:C) can reduce liver metastasis *in vivo*, the initial phenotypic changes that underpin its anti-metastatic response remain poorly described, particularly in the immunosuppressed CMS4 tumour microenvironment.

Here we characterise lineage-specific anti-metastatic responses induced by poly(I:C), including acute macrophage polarisation and a novel CMS1-like regenerative stem cell state, which drive pro-inflammatory microenvironmental changes in CRC. These insights enabled the development of tractable biomarkers that identify an “immune-warm” patient subset most likely to respond to poly(I:C), enriched for mismatch-repair proficient (pMMR), anti-inflammatory macrophages and CMS4-like features. The viral mimetic poly(I:C) offers a tailored treatment option for CMS4 tumours, by reprogramming stem cell states and activation of an innate-adaptive anti-metastatic response.

## Introduction

Colorectal cancer (CRC) is a biologically heterogenous disease. Subtyping approaches in CRC have stratified tumours into four consensus molecular subtypes (CMS)^1^ and three pathway-derived subtypes (PDS)^2^. The PDS2 group of tumours unifies the immune-rich (CMS1) and stroma-rich (CMS4) tumours into a single subtype. Immune-rich CMS1 tumours have the most favourable prognosis in terms of relapse-free survival in the localised disease setting. Furthermore, CMS1 is enriched for tumours with microsatellite instability (MSI) and deficiency in mismatch-repair (dMMR), molecular tumour types that display a strong clinical response to immune checkpoint inhibition^3^. In contrast, stroma-rich CMS4 have a poor prognosis in localised disease and represent a subtype that derives limited response to both chemotherapy and immunotherapy^4,5^.

Given the lack of response to current treatment options in these tumours, our previous work set out to identify new therapeutic options, based on the biology underpinning relapse within stroma-rich/CMS4 stage II/III tumours, where we identified two transcriptionally distinct prognostic subgroups within this otherwise uniform histological group of tumours^6^. Molecular characterisation of these two groups revealed that elevated interferon signalling, viral-related innate response and activation of antigen processing and presentation (APP) was associated with reduced relapse rates. Furthermore, we demonstrated that Polyinosinic:polycytidylic acid (poly(I:C)), a synthetic analogue of double-stranded RNA (dsRNA), could activate these phenotypic responses in *Kras*^G12D/+^ *Trp53*^fl/fl^ *Notch1*^Tg/+^ (KPN) tumours, resulting in a significant reduction of liver metastasis in this *in vivo* model of stroma-rich CMS4 CRC^7^.

Adding to our existing data supporting the therapeutic value of poly(I:C) in these stroma-rich models, here we characterise the lineage-specific mechanisms underpinning this efficacy and develop tractable biomarkers of this response, based on a comprehensive characterisation of the downstream signalling and phenotypes activated in individual cell types. Using this insight, we provide improved understanding of the processes occurring in the complex CRC tumour microenvironment during this anti-viral response and add direct evidence for the presence and extent of these characteristics in human tumours and tumour-infiltrating macrophages. These data also provide a strong rationale for future clinical translation of this therapeutic option in localised CRC.

## Results

### Poly(I:C) treatment initiates an anti-metastatic adaptive immune response

Characterisation of the responses underpinning reduced liver metastasis in KPN models following poly(I:C) treatment were initially focussed on remaining liver metastasis tissue, using flow cytometry and immunohistochemistry for the adaptive T cell response alongside digital pathology assessments of tumour burden using QuPath [Figure 1A]. Analyses of both CD8+ T cell abundance from flow cytometry data compared to tumour burden in matched mice, reveal a significant reduction in liver tumour burden in the poly(I:C) treated group relative to control (Wilcoxon rank sum, p value = 0.009), a significant increase of the CD8+ T cells relative to control (Wilcoxon rank sum, p value = 0.004) and an overall significant negative correlation between the CD8+ T cells and the liver tumour burden percentage (r=- 0.79, p value = 0.0042) [Figure 1B]. In addition to this strong correlation between cytotoxic T cells and reduced liver tumour burden, CD8+ immunohistochemistry staining in the liver tissue highlights the extensive intraepithelial CD8+ T cell infiltration within remaining metastatic lesions within the poly(I:C) treated samples [Figure 1C, Supplementary Figure 1A]. Taken together, these data highlight the anti-metastatic adaptive immune response that is driven by poly(I:C).

**Figure 1:**
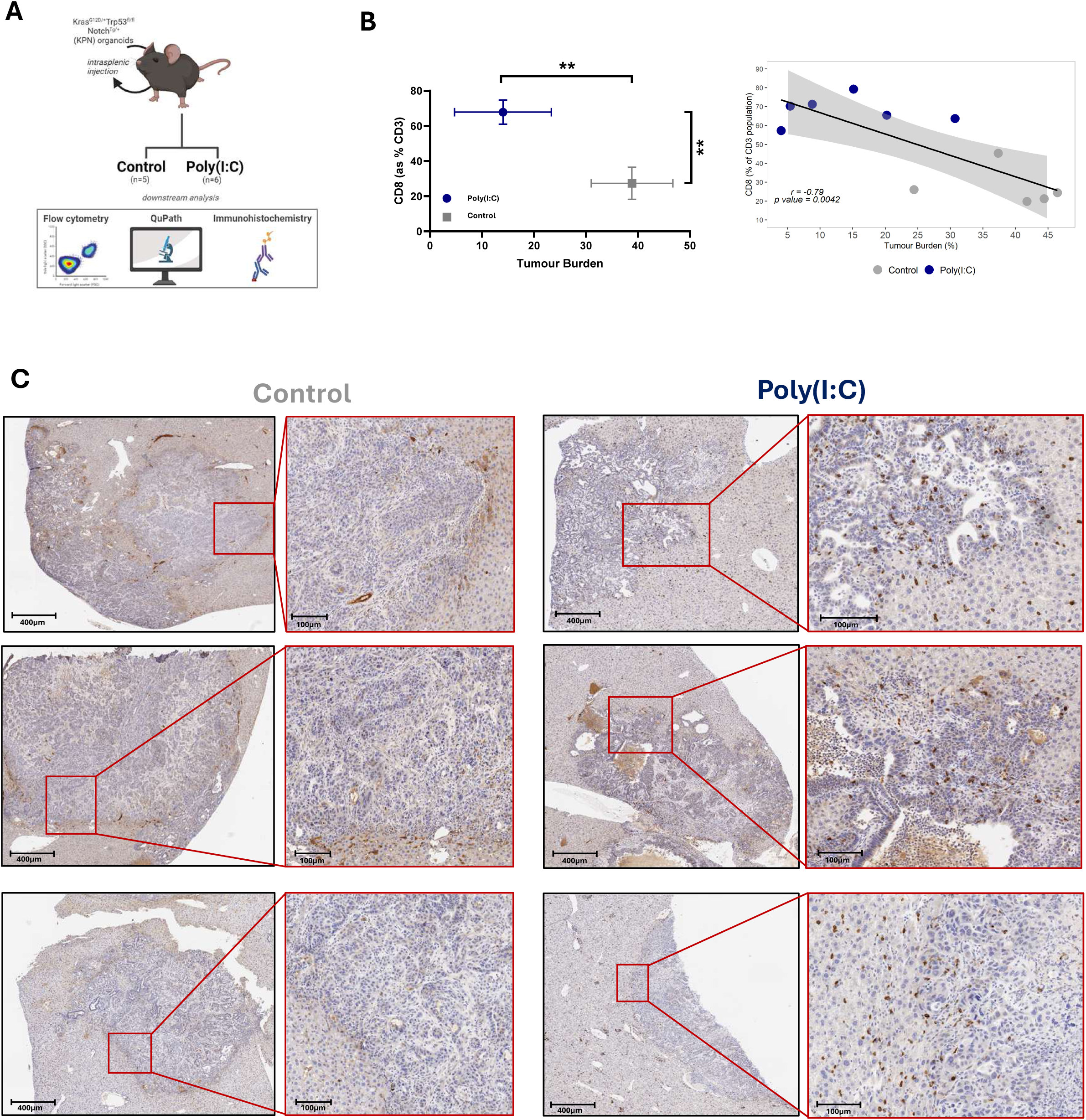
Poly(I:C) treatment induces an adaptive immune response at liver metastasis. **(A)** Schematic depicting previous *in vivo* experiment. Kras^G12D/+^Trp53^fl/fl^Notch^Tg/+^(KPN) organoids were injected using an intrasplenic model into C57BL/6 mice, which were then stratified into either a control (n=5) or poly(I:C)-treated (n=6) group. The downstream analysis of the liver metastasis involved flow cytometry, immunohistochemistry and digital pathology using QuPath. **(B)** The mean of each variable (CD8 T cell abundance (% of the CD3+ population) from flow cytometry and tumour burden quantified using QuPath) for both groups (control and poly(I:C)-treated) (Error bars indicative of standard deviation of the means), with Wilcoxon rank sum test performed for each comparison (left). Pearson’s correlation between CD8+ T cells (% of the CD3+ population from flow cytometry) and the liver tumour burden percentage (right). **(C)** Immunohistochemistry of CD8+ cells within the liver metastasis from both control and poly(I:C) treated groups. (Wilcoxon rank sum carried out by ggpubr: ns: p > 0.05, *: p <= 0.05, **: p <= 0.01, ***: p <= 0.001, ****: p <= 0.0001).

### Poly(I:C) induces distinct in vitro responses in cancer epithelial cells and THP-1-derived macrophages

While these findings highlight the anti-metastatic efficacy that poly(I:C) can achieve, they do not give us any insights into the lineage-specific mechanisms underpinning these responses, particularly those earlier in the immune activation cascade. To detangle these events, within both the tumour epithelial and innate microenvironment lineages, we initially characterised the poly(I:C) response in CRC epithelial cells (HCT116) and PMA-exposed THP-1-derived macrophages (0.04 ug/ml – 25 ug/ml) [Figure 2A]. Assessments at 24 and 48 hours reveal that while the viability of THP-1-derived macrophages remained unaffected, there is a significant (t-test, p values < 0.05) increase in cell death in HCT116 cells at all doses >0.2 ug/ml [Figure 2B, Supplementary Figure 2A-D]. Death induced by poly(I:C) is also observed in GP5D epithelial cells, however the viability of several other CRC cell lines, namely HT29, SW620 and Colo-205, is unaffected [Supplementary Figure 2E-F]. Poly(I:C) treatment results in a significant activation of both caspase-8 (7.2x fold change, t-test p value= 0.0008) and caspase-3/7 (13.4x fold change, t-test p value= 0.0014) in HCT116 cells, but not in THP-1-derived macrophages (t-test p value= 0.79 & p = 0.49 respectively) [Supplementary Figure 2G]. A significant increase in caspase-8 (1.1x fold change, t-test p=0.029) but not caspase-3/7 (1.1x fold change, t-test p=0.2) activity is observed in GP5D cells [Supplementary Figure 2H]. In line with this, caspase-8 cleaved fragments are detected in both HCT116 and GP5D cell lines when incubated with 5 ug/ml of poly(I:C), but not in poly(I:C)-treated THP-1-derived macrophages [Supplementary Figure 2I].

**Figure 2:**
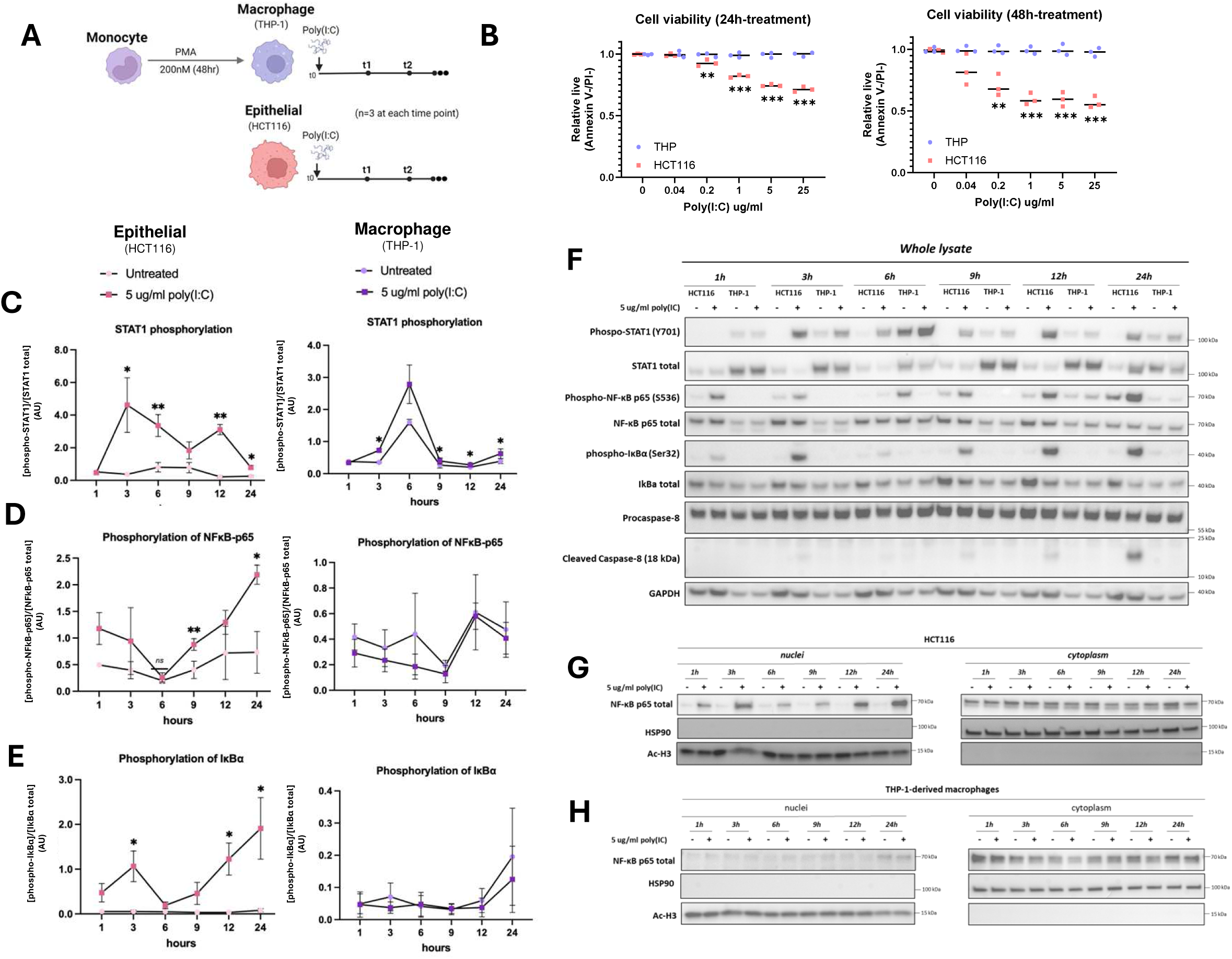
Cell lineage specific response to poly(I:C). **(A)** Schematic of the experimental outline of the response to poly(I:C). **(B)** Quantification of cell viability using AnnexinV/PI-stained cells being assessed by flow cytometry at 24hrs and 48hrs (n=3 for each cell line at each concentration). **(C-F)** Whole lysate western blots of **(C)** phosphorylated (phospho) STAT1/STAT1 total, **(D)** phospho NFkB(p65)/ NFkB(p65) total and **(E)** phospho IkBα/ IkBα total across 6 time points with the absence or addition of poly(I:C). (n=3 for each cell line at each time point). **(G)** Nuclear and cytoplasmic NFkB (p65) in the epithelial (HCT116) and **(H)** THP-1 macrophages across 6 time points with the absence or addition of poly(I:C). (Student t-test carried out on Prism: ns: p > 0.05, *: p <= 0.05, **: p <= 0.01, ***: p <= 0.001, ****: p <= 0.0001). Error bars = 1 standard deviation.

In addition to viability changes, we observe activated signalling of viral-response related transcription factors, namely STAT1 and NFkB, in HCT116 cells following poly(I:C) treatment, as evidenced by increased p65 and IkBα phosphorylation levels and phosphorylation of STAT1 [Figure 2C-F]. Activation of NFkB pathway by poly(I:C) is further confirmed by assessment of nuclear localisation of p65 in HCT116 cells [Figure 2G]. In contrast, in THP-1-derived macrophages, only STAT1 activation is observed, with an absence of p65 and IkBα phosphorylation observed in either whole cell lysates or nuclear extractions [Figure 2C-F, 2H].

### Lineage-specific poly(I:C) responses drive several overlapping and distinct phenotypes

To unbiasedly identify downstream signalling responses following poly(I:C) treatment, we transcriptionally profiled both epithelial (HCT116) and macrophage (THP-1) cell lines in the absence and presence of 5 ug/ml poly(I:C) after 24 hours [Figure 3A]. Analysis of the biological Hallmark gene sets using gene set enrichment analysis (GSEA) reveals that poly(I:C) treatment in these epithelial cells leads to a significant enrichment in interferon alpha and gamma response, tumour necrosis factor-alpha (TNF-alpha) via NFkB signalling, as well as immune pathways including inflammatory response and complement [Supplementary Figure 3A]. In addition to these epithelial-specific responses, both epithelial and macrophage treated cell lines have a significant enrichment in interferon (IFN)-alpha and IFN-gamma response (adjusted p value <2.6e-45, NES >3 across all analysis) [Figure 3B]. To test if the differences between the poly(I:C) responses were due to a difference in the baseline transcriptional signalling prior to treatment in these two cell lines, we next assessed these interferon-related signatures using single sample gene set enrichment analysis (ssGSEA). This comparison confirms the significant enrichment of interferon-pathways upon poly(I:C) treatment, but also reveals that untreated macrophages have a significantly higher baseline level of both interferon response pathways than untreated epithelial cells (Wilcoxon rank sum test, * = p value < 0.05) [Figure 3C].

**Figure 3:**
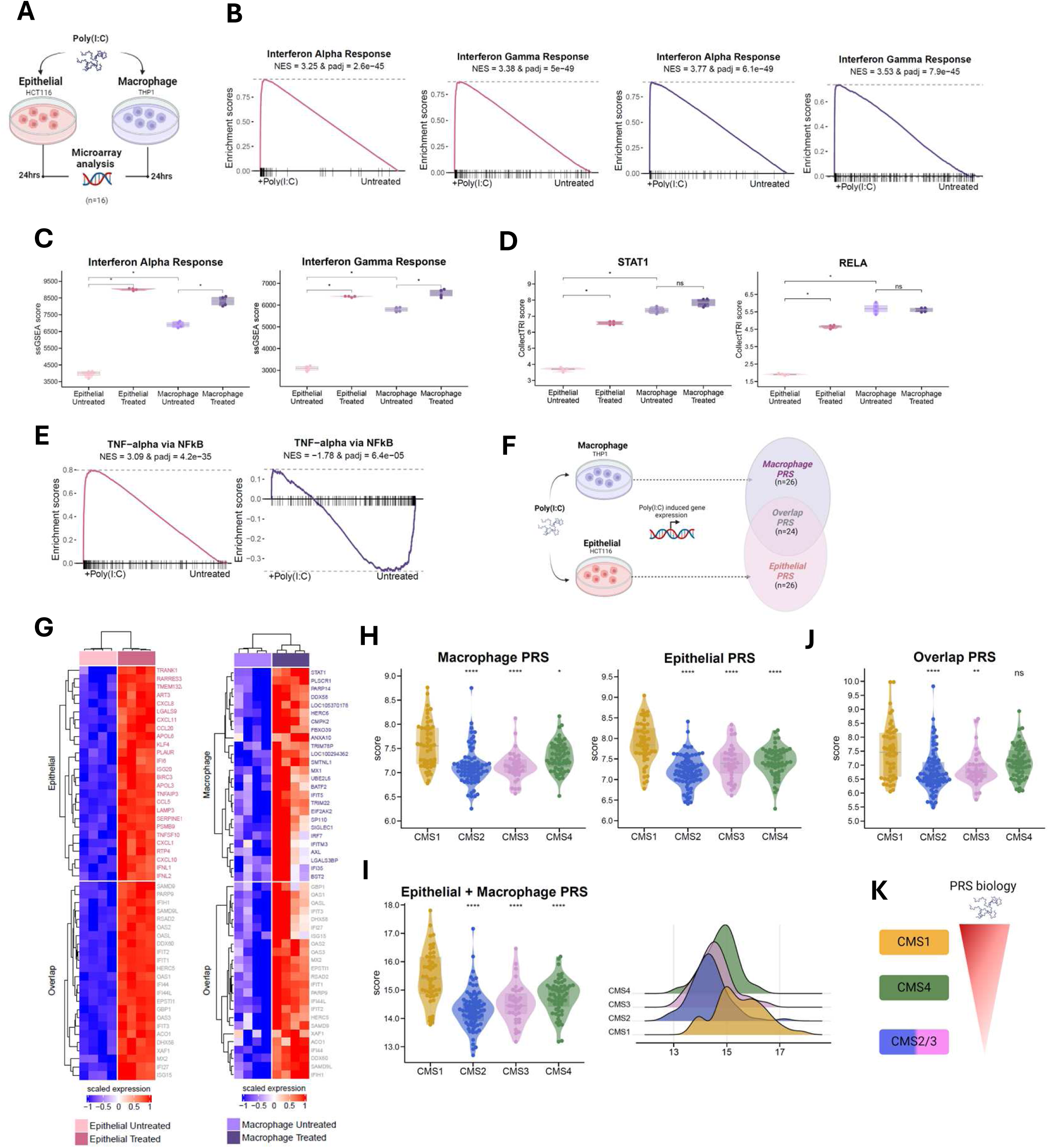
Transcriptional analysis of *in vitro* model. **(A)** Outline of experimental design of the microarray analysis. (n=16 samples, n=4 in each cell line for each condition). **(B)** Enrichment plots of the gene set enrichment analysis (GSEA) results of Interferon-alpha response and Interferon-gamma response (epithelial;pink and macrophage; purple). **(C)** Single sample gene set enrichment analysis (ssGSEA), of these same interferon gene sets. **(D)** Transcription factor activity scores of STAT1 and RELA. **(E)** Enrichment plots of the GSEA results of TNF-alpha via NFkB for epithelial (pink) and macrophage (purple). **(F)** Schematic illustrating the development of the poly(I:C) response signatures (PRS), macrophage (n=26 genes), epithelial (n=26 genes) and overlap (n=24 genes). **(G)** Expression of the PRS genes across the treated and untreated cell lines. **(H)** Macrophage PRS (left) and Epithelial PRS (right), **(I)** Macrophage PRS + Epithelial PRS and **(J)** Overlap PRS was compared across the consensus molecular subtypes (CMS) in transcriptional profiles from a stage II/III colon cancer cohort (GSE39582), (n=258; CMS1 = 49, CMS2 = 75, CMS3 = 35, CMS4 = 58, unknown = 41) (Wilcoxon rank sum test, with CMS1 as the reference group). **(K)** Schematic highlighting the PRS biology highest in CMS1, followed by CMS4 and then CMS2/3. (Wilcoxon rank sum carried out by ggpubr: ns: p > 0.05, *: p <= 0.05, **: p <= 0.01, ***: p <= 0.001, ****: p <= 0.0001).

Transcription factor activity assessments [Figure 3D] confirmed a significant activation of STAT1 in the treated epithelial cells (Wilcoxon rank sum test, p value < 0.05) [Figure 3D]. Further alignment with the initial analysis highlights that the NFkB pathway is uniquely activated in the treated epithelial cells, shown through activation of RELA [Figure 3D], as well as significant enrichment of the TNF-alpha via NFkB pathway [Figure 3E]. These data so far have revealed how treatment with poly(I:C) can stimulate both overlapping and unique transcriptional responses in epithelial and macrophage cell lineages.

### Poly(I:C) upregulates transcriptional signalling that resembles a CMS1-like tumour state

To identify these overlapping and lineage-specific responses to poly(I:C), we selected the top n=50 upregulated genes in each cell line, relative to their control, following poly(I:C) treatment [Figure 3F]. In line with the overlapping biology identified earlier, many of these upregulated genes are shared across the two cell line models, resulting in three final poly(I:C) response signatures (PRS); epithelial (n=26 genes), macrophage (n=26 genes) and overlap (n=24 genes) [Figure 3F-G]. When these signatures were assessed in transcriptional profiles from stage II/III colon tumours (GSE39582) (n=258), the PRS macrophage and epithelial signatures, either individually [Figure 3H] or in combination [Figure 3I], alongside the overlap PRS [Figure 3J], are found to be significantly higher in CMS1 tumours followed by CMS4, with CMS2/3 tumours having the lowest expression (Wilcoxon rank sum test, significance = p value < 0.05). Overall, these data indicate that the biology induced by poly(I:C) in both epithelial and macrophage lineages, which we know results in an anti-metastatic cytotoxic response *in vivo* [Figure 1], strongly aligns with the overall tumour biology of a CMS1 tumour [Figure 3K].

### The biology induced by poly(I:C) results in polarisation of macrophage populations towards a pro-inflammatory, anti-tumour phenotype

An assessment of macrophage abundance indicates that there is no significant difference between CMS1 and CMS4 tumours for these innate lineages overall (Wilcoxon rank sum test, p value = 0.9551) [Figure 4A]. However, when these same tumours are assessed for a previously-defined transcriptional signature^8^ associated specifically with an pro-inflammatory anti-tumorigenic M1-like phenotype, CMS1 tumours have a significantly higher score compared to all other groups, including CMS4 tumours, when used directly [Supplementary Figure 4A] or as a proportion of all macrophages (Wilcoxon rank sum test, p value = 0.0006) [Figure 4B]. Furthermore, using this pro-inflammatory macrophage transcriptional signature, alongside STAT1 and STAT2 as additional markers of M1 macrophages, we observe that these are all significantly increased in the poly(I:C) treated macrophages relative to untreated (Wilcoxon rank sum test, M1-like signature p value = 0.029, STAT1/2 p value = 0.00016, t-test, CD80 p value < 0.05) [Figure 4C, Supplementary Figure 4B], further supporting the therapeutic value of poly(I:C) in this setting. This contrasts with the significant elevation of anti-inflammatory M2-like signalling in CMS4 tumours [Supplementary Figure 4C]. Alongside this increased expression, we also observe elevated nuclear staining of both STAT1 and STAT2 at 6 and 24 hours post poly(I:C) treatment in THP-1 macrophages [Figure 4D].

**Figure 4:**
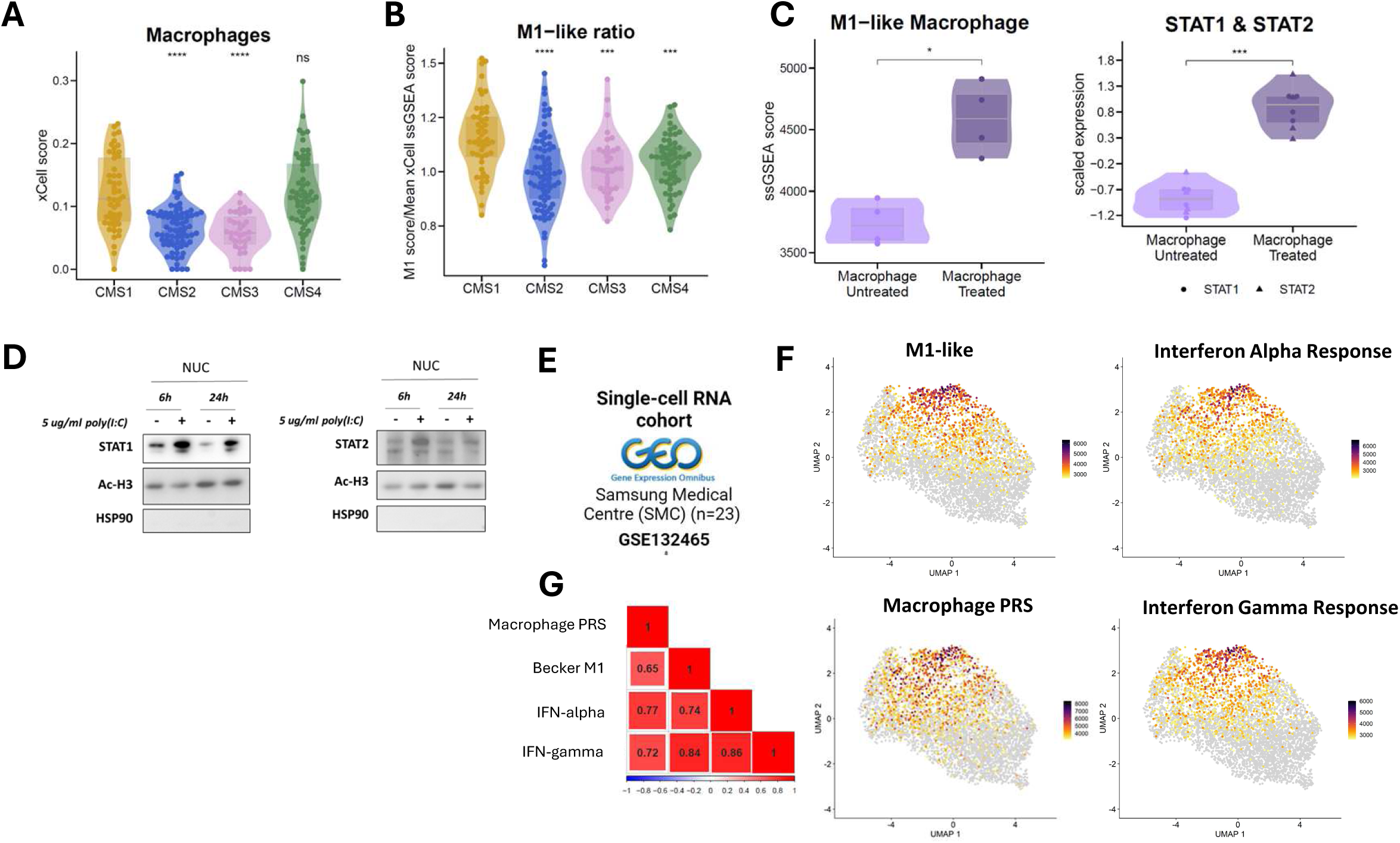
Poly(I:C) induces an M1-like phenotype in macrophages which is encapsulated within the macrophage PRS. **(A)** xCell macrophage score across the CMS’s and **(B)** M1-like transcriptional signature single sample gene set enrichment analysis (ssGSEA) score, divided by the mean ssGSEA score of xCell macrophage gene lists (GSE39582) (Wilcoxon rank sum test, with CMS1 as reference group). **(C)** ssGSEA of M1-like signature (left) and scaled expression of M1 surrogate genes (STAT1 and STAT2) in macrophages treated and untreated with poly(I:C) (right). **(D)** STAT1 and STAT2 nuclear extraction within treated and untreated macrophages at both 6- and 24-hour time points. **(E)** Schematic of single cell RNA sequencing cohort that was utilised (GSE132465) (n=23). **(F)** Myeloid subset of cells (n=5586) that were scored with M1-like transcriptional signature, Macrophage PRS, Interferon Alpha and Interferon gamma response. **(G)** Spearman’s correlation of the scores from **(F)**, (blue to red for negative to positive correlation). (Wilcoxon rank sum carried out by ggpubr: ns: p > 0.05, *: p <= 0.05, **: p <= 0.01, ***: p <= 0.001, ****: p <= 0.0001).

To validate these findings in more granular data, we utilised a single cell RNA-sequencing cohort derived from n=23 CRC tumour samples (GSE132465) and selected the macrophage population (n=5586 cells) for further analysis [Figure 4E]. Applying the same transcriptional signatures used earlier, in both the bulk tumour cohort and *in vitro* models, we observe that the subset of macrophage cells that displays the highest expression levels for the M1-like signature also has the highest expression levels for macrophage PRS, interferon alpha and gamma; capturing the biologies and phenotypes that can be induced by treatment with poly(I:C) [Figure 4F,G]. Stratification of the macrophage cells into pro-inflammatory M1-like high and low subsets, again highlights the significant overlap between pro-inflammatory macrophages and the biology induced by poly(I:C) in these lineages (Wilcoxon rank sum test, p value <2e-16) [Supplementary Figure 4D]. In the bulk tumour cohort, we observe that while CMS2/3 tumours are uniformly low for this pro-inflammatory signature, CMS1 tumour appear to have a bimodal distribution, where CMS1 tumours with low levels of pro-inflammatory signalling display similar signalling to that of CMS4 tumours [Supplementary Figure 4E]. Further stratification of CMS1 based on M1 signalling, into high and low subsets, again confirms that the phenotypes induced during a poly(I:C) response are significantly associated with the strongest M1-like signalling within CMS1 tumours, rather than supressed CMS4 tumours, or immune low CMS2/3 (Wilcoxon rank sum test, p value = 6.842e-09) [Supplementary Figure 4F]. Taken together, these findings confirm that the response to poly(I:C) results in a polarisation of macrophages towards a proinflammatory activated phenotype in both our *in vitro* lineages and a strong alignment between a poly(I:C) response and M1-like macrophages in bulk and single cell tumour data.

### Poly(I:C) induces a CMS1-like regenerative stem state

We next set out to identify epithelial phenotype associations with poly(I:C) response, with particular focus on stem populations, using the intestinal stem cell index^9^. In our *in vitro* models, we observe a significant increase in the stem cell index in poly(I:C) treated cells compared to untreated, indicating a shift away from a canonical crypt-base columnar (CBC) stem cell phenotype towards a more regenerative stem cell (RSC) state (Wilcoxon rank sum test, p value = 0.02857) [Figure 5A]. When we measure either the stem cell index or RSC, compared to epithelial PRS scores, we find a significant positive correlation (stem cell index; r = 0.58, p value = 2.8e-21, RSC; r = 0.6, p value = 9.7e-23) [Figure 5B]. However, despite these clear statistical correlations, when we assess RSC across the CMS subtypes we find no measurable difference in this regenerative phenotype between CMS1 and CMS4 tumours, although both are significantly elevated compared to CMS2/3 tumours [Figure 5C, Supplementary Figure 5A].

**Figure 5:**
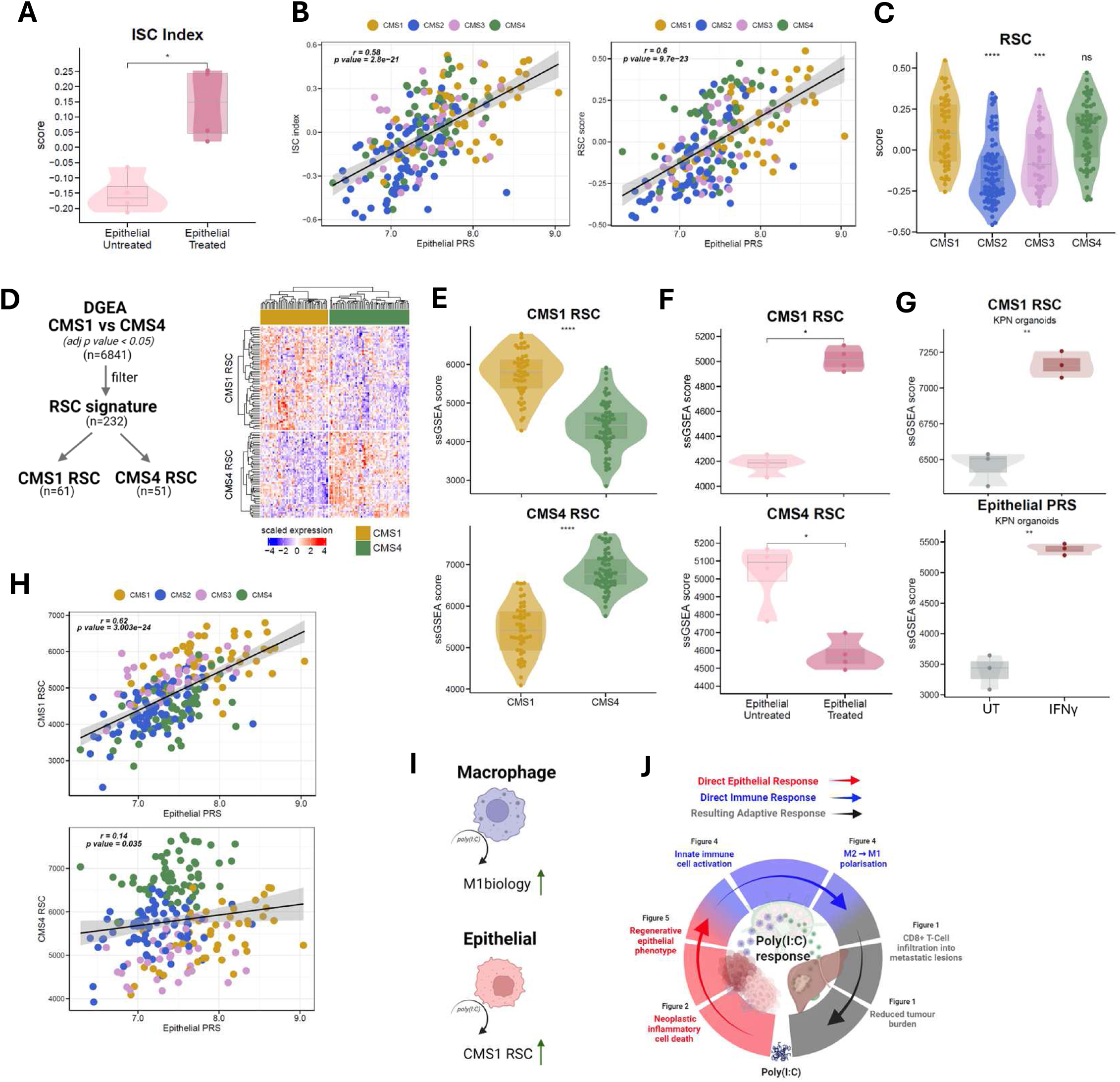
Poly(I:C) induces a CMS1-like regenerative stem state. **(A)** Intestinal stem cell index between the poly(I:C) treated and untreated epithelial cells. **(B)** Pearson’s correlation of the stem cell index scores (left) and regenerative stem cell (RSC) score with epithelial PRS score (right) (GSE39582) (n=258). **(C)** RSC score across the CMS’s (Wilcoxon rank sum, with CMS1 as reference group). **(D)** Flow diagram of the creation of the CMS1 RSC and CMS4 RSC gene signatures (left) and their visualisation (right). **(E)** Single sample gene set enrichment analysis (ssGSEA) scores of the CMS1 RSC (top) and CMS4 RSC (bottom) gene signatures between CMS1 and CMS4 samples. **(F)** ssGSEA scores of the CMS1 RSC and CMS4 RSC gene signatures in the poly(I:C) treated and untreated epithelial cells. **(G)** ssGSEA scores of CMS1 RSC (top) and Epithelial PRS (bottom) in KPN Organoids treated with interferon-gamma (IFNγ) (E-MTAB-11769) (t-test). **(H)** Pearson’s correlation of the CMS1 RSC (top) and CMS4 RSC (bottom) ssGSEA scores with epithelial PRS. **(I)** Schematic illustrating that poly(I:C) induces a M1-like phenotype in macrophages and a CMS1-like regenerative state in epithelial cells. **(J)** Steps of the cancer immunity cycle that are part of the poly(I:C) response. Each step has been shown in a corresponding figure. (Wilcoxon rank sum carried out by ggpubr: ns: p > 0.05, *: p <= 0.05, **: p <= 0.01, ***: p <= 0.001, ****: p <= 0.0001).

As this regenerative state has been associated with both a stromal and inflammatory microenvironment, we attempted to segregate this RSC score into genes associated with a CMS1 RSC, with a more immune active inflammatory phenotype, and RSC genes associated with CMS4, with a more stromal immune supressed phenotype. To do this, we performed a differential gene expression analysis between CMS1 and CMS4 and overlapped the significant genes (adjusted p value < 0.05, n=6841 genes) with the RSC signature (n=232), which resulted in two signatures referred to as CMS1 RSC (n=61/232) and CMS4 RSC (n=51/232) [Figure 5D]. Using these new signatures, we confirm that as expected the CMS1 RSC is significantly higher in CMS1 tumours relative to CMS4 tumours (Wilcoxon rank sum test, p value = 1.5e-14), while CMS4 RSC is significantly higher in CMS4 tumours relative to CMS1 tumours (Wilcoxon rank sum test, p value = < 2.2e-16) [Figure 5E]. Most importantly, however, we now show that these distinct regenerative states are regulated in entirely opposing directions during response to poly(I:C), with significant upregulation of the CMS1 RSC genes in the poly(I:C) treated epithelial cells, with concurrent significant downregulation of the CMS4 RSC signature (Wilcoxon rank sum test, both p values < 0.05) [Figure 5F, Supplementary Figure 5B]. Furthermore, KPN organoids treated with interferon gamma (IFNγ) (E-MTAB-11769) were significantly enriched for both the CMS1 RSC (t-test, p value = 0.0017) and epithelial PRS (t-test, p value = 0.0034) (Figure 5G). However, when KPN organoids were treated with TGF-β (E-MTAB-11784), the CMS1 RSC ssGSEA score was significantly lower than that of the untreated organoids (t-test, p value = 0.041) (Supplementary Figure 5C). In addition, we also see that using these refined regenerative stem cell signatures, it is only the CMS1 RSC that displays a strong positive correlation with the epithelial PRS (r = 0.62, p value = 3.003e-24), while the CMS4 RSC has a very weak correlation (r = 0.14, p value = 0.035) [Figure 5H].

Overall, by stratifying the broad regenerative stem cell signature into two distinct phenotypes, based on CMS1 v CMS4 human tumour biology, alongside the use of M1-like markers in human tumour macrophages, we highlight that poly(I:C) treatment drives a strong phenotypic change in both epithelial and immune tumour lineages, through a combined elevation of a CMS1-like RSC and pro-inflammatory macrophage state, while supressing the CMS4-like RSC and supressed macrophage state [Figure 5I]. Coupled to this regenerative shift, given the direct inflammatory cell death induced in some of our epithelial models [Figure 2] and the clear cytotoxic response we see in the livers of metastatic mouse models [Figure 1], these data highlight how the overall responses to poly(I:C), both direct and indirect, can be used to overcome an immunosuppressed tumour microenvironment by triggering a number of critical cascades in parallel within the anti-cancer immunity cycle [Figure 5J].

### Poly(I:C) response aligns with biological characteristics of immune-hot CMS1 tumours

To further investigate the alignment of the PRS biology with human tumour subtypes, the expression of the three signatures was used in combination to generate a PRS sum score (see methods), which enabled human colorectal tumours to be ranked along a biological gradient from high PRS to low PRS [Figure 6A]. CMS1 tumours overall display the highest poly(I:C) response biology, with the samples lowest being predominantly classified as CMS2/3 [Figure 6A-B]. However, while this is clearly a continuum of biological signalling, to provide granularity on the relationship between PRS biology and the binary subtype classifications, tumours samples were split into a Hot (top 25%), Warm (intermediate 25%) and Cold (lowest 50%) subgroups based on their expression of PRS scores [Figure 6C-D]. dMMR tumours are associated with the Hot and lesser extent Warm PRS groups, compared to Cold [Figure 6D, Supplementary Figure 6A]. Using PHATE to visualise these data, we confirm that the highest expression of poly(I:C) response biology is in the Hot group, followed by Warm and then Cold tumours [Figure 6E] (left to right). Tumours within the Hot subgroup are predominantly CMS1, whereas CMS4 tumours predominantly classified as Warm, with Cold tumours mostly classified as CMS2/3 [Figure 6F]. In line with the previously defined association between PDS and CMS^2^, we find that more than 85% of Hot/Warm tumours are PDS2 [Figure 6G, Supplementary Figure 6B]. Furthermore, the Hot tumour group is significantly elevated for signatures of innate immune response, antigen processing and presentation (APP) and immune signalling in human tumours [Figure 6H-I, Supplementary Figure 6C]. High expression of all PRS scores was associated with a significantly lower relapse rates in both the GSE39582 (n=258) (HR = 3.802, p value = 0.0054) and an independent stage II colon cancer cohort (E-MTAB-863) (n=215) (HR = 2.252, p value = 0.00055) [Figure 6 J-K] [Supplementary Figure 6D-E].

**Figure 6:**
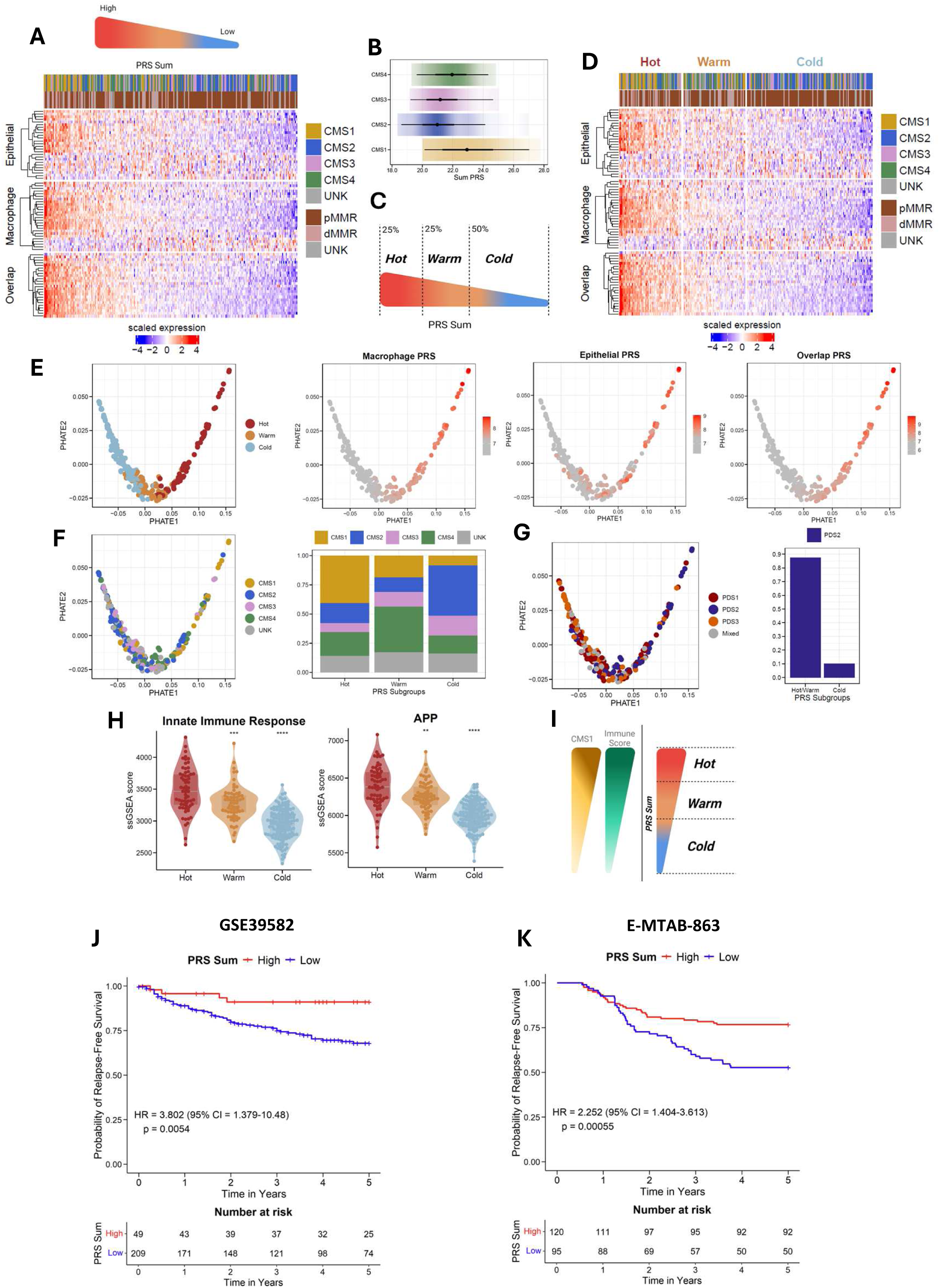
Poly(I:C) response aligns with biological characteristics of immune-hot CMS1 tumours**. (A)** Transcriptional samples ranked based on their expression of the poly(I:C) response signature (PRS) genes, with annotations of their CMS classification and mismatch repair status (MMR) (p; proficient and d; deficient) (GSE39582) (n=258). **(B)** The sum of the three PRS scores (macrophage, epithelial and overlap) (PRS sum) across the CMS’s. **(C)** Schematic depicting the stratification of the samples into three PRS subgroups; Hot, Warm and Cold and **(D)** their visualisation. **(E)** PHATE visualisation of the PRS subgroups, Macrophage PRS, Epithelial PRS and Overlap PRS (left to right). **(F)** PHATE visualisation of CMS classification (left) and proportion of each CMS classification within each PRS subgroup (right). **(G)** Pathway derived subtypes (PDS) classification visualised using PHATE (left), with the percentage of Hot/Warm and Cold samples that were classified as PDS2 (right). **(H)** Single sample gene set enrichment analysis scores (ssGSEA) of innate immune response (left) and antigen processing and presentation (right) across the PRS subgroups (Wilcoxon rank sum test, with Hot subgroup as the reference group). **(I)** Schematic depicting the alignment of CMS1-like biology and immune score with the PRS subgroups. **(J)** Kaplan-Meier survival estimates of GSE39582 (n=258) and an independent stage II colon cancer cohort (E-MTAB-863) (n=215) calculated using the PRS sum score. (Wilcoxon rank sum carried out by ggpubr: ns: p > 0.05, *: p <= 0.05, **: p <= 0.01, ***: p <= 0.001, ****: p <= 0.0001).

## Discussion

In this study, we set out to characterise a series of key downstream signalling and phenotypic cascades induced following the anti-viral response triggered by poly(I:C) treatment, and to shed new light on the key processes underlying its anti-metastatic efficacy in pre-clinical models. The biology induced in each lineage type was summarised into a series of new biology-driven gene signatures, referred to as poly(I:C) response signatures (PRS). The use of PRS scores enabled us to test the human tumour relevance of this biological response across independent cohorts, including single cell RNAseq and bulk stage II/III CRC. Using these biology-driven biomarkers, data presented here highlights how tumours that are classified as either CMS2 or CMS3 are overwhelmingly biased towards developing an overall cold, immune excluded, microenvironment. In contrast, tumours classified as PDS2, which is a subtype enriched for both CMS1 and CMS4 tumours, were predominantly set on a developmental path that will bias them towards establishing either a hot or warm immune microenvironment. Characterisation of the hot tumour group revealed a number of key immune and epithelial features associated with a proinflammatory (M1-like) phenotype, inflammatory cell death and a new CMS1-like regenerative stem cell state. Although CMS4 tumours have many CMS1-like characteristics, their immunosuppressive tumour microenvironment results in its classification as PRS warm, with characteristics that contrast with the features of the hot PRS group, namely an anti-inflammatory (M2-like) phenotype and a new CMS4-like regenerative state.

These hot and warm phenotypic tumour landscapes observed at diagnosis, if left unperturbed, would progress to their well-characterised opposing clinical outcomes associated with these distinct molecular subtypes. However, the cells within these tumour ecosystems still retain many features associated with cancer cell plasticity and immune cell polarisation, which can be mediated by acute external stimuli. Our data strongly support the therapeutic value of using the clinically safe compound poly(I:C) to phenotypically shift the immunosuppressive immune and epithelial phenotypes within this warm group towards those of a pro-inflammatory and CMS1-like hot phenotypic state, in a way that triggers a highly efficacious anti-metastatic adaptive immune response [Figure 7].

**Figure 7:**
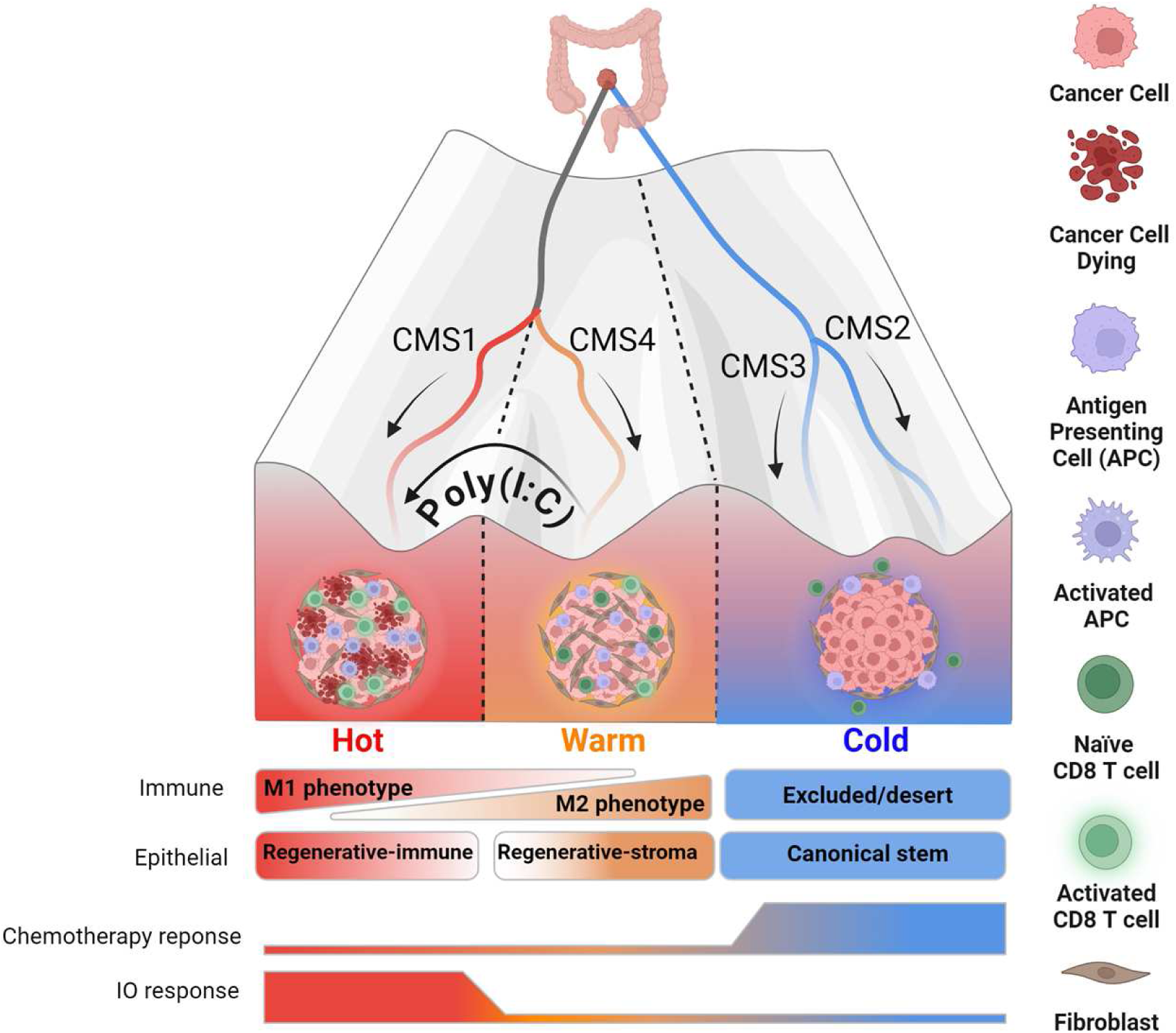
Using the poly(I:C) response biology, we identify a PRS Hot group enriched for PDS2/CMS1 tumours, with a pro-inflammatory and CMS1-like regenerative phenotype. Aligning with our understanding of this immune-active CMS1 and dMMR enrichment, this Hot PRS subgroup would be most responsive to immunotherapy. The PRS cold group is enriched for epithelial-rich CMS2/3 tumours, tumour subtypes that have been shown to derive some response to standard of care chemotherapy. The PRS Warm group was enriched for PDS2/CMS4 tumours, with these stroma-rich characteristics associated with a lack response to many therapeutics, including chemotherapy and immunotherapy. PRS Warm tumours are enriched for pro-tumour M2 macrophages along with a CMS4-like regenerative stem phenotype. Treatment of poly(I:C) can increase all of these important biologies, which provide a therapeutic opportunity to shift the phenotypic landscape of Warm tumours so that they resemble the phenotypic landscape of a Hot tumour and the associated anti-metastatic adaptive properties.

Our approach revealed that the phenotypes stimulated by poly(I:C) were sufficient to stratify tumour samples along an immune and epithelial phenotypic landscape, from hot (immune-rich/CMS1-dMMR enriched), to warm (immune-intermediate, CMS4 enriched) and cold (immune-devoid/CMS2-3 enriched). The cold tumours were characterised by an epithelial-rich phenotype, which are known to be responsive to current chemotherapies.^10^ In contrast, the hot subgroup was enriched for the clinical feature of dMMR, a biomarker that is associated with tumours that display the highest responses to immune checkpoint blockade therapy. Therapeutic interventions in CRC with ICB’s in the neoadjuvant setting has shown remarkable success rates, with 100% of dMMR tumours displaying clinical response^11^. Interferon signalling can induce the activation of an array of different transcription factors, among which STAT1 plays an essential role in the regulation of the expression of many interferon-stimulated genes (ISGs)^12,13^. Interferon-induced STAT1 and STAT2 signalling has been associated with the activation of anti-tumour properties in antigen presenting cells^14^ and promotion of APP via expression of lymphocyte-stimulating glycoproteins such as CD80^15^. These findings align with the M1-like phenotype induced by poly(I:C) in this study, represented by enhanced APP and STAT1 protein.

While these hot and cold groups have been described in the past, our study provides more precise evidence for the presence of a warm subgroup, one which is enriched for the poor-prognostic stroma-rich CMS4 subtype and currently derives limited clinical benefit from many therapeutic interventions^4,5,16^. The approach of using a ‘CMS4-switch-therapy’, has been investigated previously^17^, whereby a selective pressure is applied to skew the biology into a more favourable CMS-phenotype. In line with this hypothesis, our *in vitro*, *in vivo* and *in silico* analyses indicate the therapeutic potential for neoadjuvant treatment of immunosuppressed pMMR tumours with poly(I:C), to enable triggering of inflammatory intrinsic cell death, an M1-like polarisation of macrophages via a viral response, and a phenotypic shift towards a microenvironmental landscape that resembles a prognostically-favourable CMS1.

In summary, our findings outline how the lineage-specific poly(I:C) responses combine to reactivate multiple steps within the cancer immunity cycle, triggering several important inflammatory signalling cascades that are observed within heterogeneous bulk tumour tissue and single cell data. Despite the cell-type specificity of these responses, the poly(I:C) response activates multiple levels of a highly coordinated anti-tumour process, which appears to culminate in a cytotoxic activation. As such, these data strongly support the use of poly(I:C) as a therapeutic opportunity in localised immunosuppressed pMMR CRC prior to surgery, to enable a widespread triggering of an innate-adaptive activation that ultimately unleashes a cytotoxic response in micro-metastatic lesions.

## Materials and methods

### Cell culture

THP-1 (RRID: CVCL_0006), HCT116 (RRID: CVCL_0291), Colo-205 (RRID: CVCL_0218), GP5D (RRID: CVCL_1235), SW620 (RRID: CVCL_0547) and HT29 (RRID: CVCL_0320) were used. HCT116 cell lines were cultured in Dulbecco’s modified Eagle’s medium (DMEM) supplemented with 10% FBS, 50 U/mL penicillin, 0.1 mg/mL streptomycin, 2 mM l-glutamine, and 1 mM sodium pyruvate (all Gibco). THP-1 cells were obtained from American Type Culture Collection (ATCC, Manassas, VA, USA) and cultured in RPMI medium (Invitrogen, Paisley, UK) with 10% heat-inactivated FBS, 50 U/mL penicillin, 0.1 mg/mL streptomycin and 0.05 mM β-mercaptoethanol (Sigma). All cells were maintained at 37 °C in a 5% CO_2_ humidified atmosphere and regularly screened for the presence of mycoplasma using the MycoAlert Mycoplasma Detection Kit (Lonza). Cell lines were cultured for no more than 20 passages following thawing. THP-1 cells were differentiated to M0 macrophages by 48 h treatment with 200 nM Phorbol 12-myristate 13-acetate (Sigma-Aldrich) in complete RPMI media. After removing the media, THP-1-derived macrophages were incubated with indicated poly(I:C) concentrations in serum-free RPMI media. Poly(I:C) treatments of HCT116 cells were also performed in DMEM serum-free media.

### Cell viability

Following 24h (and 48h) poly(I:C)-treated cells were harvested, and cell viability was analysed by trypan blue counts or by flow cytometry of Annexin V/PI-stained cells. Both, floating and adherent cells were collected for cell viability investigation. Death rate of Annexin V/PI-stained cells was assessed on the BD AccuriTM C6 flow cytometer (BD Biosciences).

### Caspase activity assay

Caspase-3/7 and -8 activities were assayed using Caspase-Glo®- 3/7 and -8 assays according to manufacturer’s instructions (Promega, Madison, WI).

### Subcellular fractionation

Nuclear and cytoplasmic fractions were obtained as previously described^18^. Briefly, the cells were lysed for 20 minutes at 4°C in “Buffer A” (10mM HEPES pH7.4, 1.5mM MgCl_2_, 10mM NaCl, 0.1% NP40, 1mM PMSF, 0.1mM TLCK, 0.1mM TPCK, 1mM NaF, 1mM Na_3_VO_4_); followed by centrifugation at 12,000g for 2 minutes; collected supernatants were then further centrifuged for 2 minutes to yield the cytoplasmic fraction. The pellets obtained from the first centrifugation were subjected to further lysis in ‘Buffer C’ (10mM HEPES pH7.4, 1.5mM MgCl_2_, 420mM NaCl, 0.1% NP40, 1mM PMSF, 0.1mM TLCK, 0.1mM TPCK, 1mM NaF, 1mM Na_3_VO_4_) in presence of Benzonase (Sigma) for 30 minutes at 4°C. Nuclear fractions were then obtained by centrifuging at 12,000g for 2 minutes the lysates and collecting the supernatant.

### Western blotting

After removing the media, cold PBS was added to the plates and the cells were scraped from the plate. Cell suspensions were then centrifuged, washed again with PBS, and the pellets were lysed in in RIPA Buffer supplemented with PhosSTOP (Roche) and Protease Inhibitor Cocktail (Roche). Equal amounts of protein (10-30 µg/well) were resolved by SDS-PAGE and transferred to nitrocellulose membranes using iBLOT (Invitrogen). Westerns were developed digitally using a FluorChem SP imaging system (Alpha Innotech) or the G:BOX ChemiXX6 gel doc system (Syngene). The following Western blot antibodies were used: Phospho NF-κB p65/RelA (S536) (RRID: AB_331284), Phospho IκBα (S32) (RRID: AB_561111), IκBα (RRID: AB_390781), STAT1 (RRID: AB_2198300), STAT2 (RRID: AB_2271323) (all Cell Signalling Technology); GAPDH (Abcam, RRID: AB_2107448), NF-κB p65 (Total RelA) (Santa Cruz Biotechnology, RRID: AB_628017),; Acetyl Histone H3 (EMD Millipore); Caspase-8 (both Enzo Life Sciences) (RRID: AB_2050949), pSTAT1 (Y701) (Invitrogen, ThermoFisher) (RRID: AB_2533113).

### RNA isolation

After removing the media, cold PBS was added to the plates and the cells were scraped from the plate. Cell suspensions were then centrifuged, and the cell pellets were re-suspended in ice-cold PBS. Total RNA was extracted from HCT116 and THP-1 cell lines using the Roche High Pure FFPE Extraction Kit (Roche Life Sciences) and amplified using the NuGen Ovation FFPE Amplification System v3 (NuGen San Carlos). The amplified product was hybridized to the Almac Diagnostics cDNA microarray-based XCEL array (Almac) and analysed using the Affymetrix Genechip 3000 7G scanner (Affymetrix). Microarray raw CEL files which were processed, and RNA normalized using the affy (1.68.0)^19^ (RRID:SCR_012835) and affyio (1.60.0)^20^ (RRID:SCR_024223) packages. The probes were collapsed to gene-level data using “MaxMean” within the WGCNA (1.7.1)^21^ (RRID:SCR_003302) package.

### *In vivo* experiment

The intrasplenic model was performed as described in our previous publication^6^, using Kras^G12D/+^, Trp53^fl/fl^ and constitutively activated NOTCH1 (KPN) organoids which underwent intrasplenic injection into C75Bl/6 mice. The organoid recipient and donor were sex and strain matched. The first treatment with poly(I:C) was at nine days following implantation (4mg/kg in saline) (n=6), with the control untreated group receiving saline vehicle control (n=6, n=5 used for tissue processing) and it was then biweekly up until day 42. A biopsy of the liver was taken for flow cytometry with the remainder fixed and used for downstream QuPath analysis (RRID:SCR_018257). For flow cytometry, detailed description of the tissue processing and gating strategies can be found in our previous publication^6^ (Supplementary methods and figures).

### Bioinformatic analysis, bulk and *in vitro*

Transcriptional profiles of stage II/III untreated colon cancer (CC) (GSE39582) (n=258) and the stage II colon cancer cohort (E-MTAB-863) (n=215) were downloaded and normalized as previously described^6^. For each PRS score, the surv_cutpoint function was used with relapse free survival (5 years) used as the optimal cutpoint variable. The function survfit and ggsurvplot were used to generate survival data and plots respectively. The packages survival (3.5.8) (RRID:SCR_021137) and survminer (0.4.9) (RRID:SCR_021094) were used. The hazard ratios were calculated on these newly stratified groups using the coxph function. R version 4.0.5 was used with seed set to 121 to carry out normalization, classification calls and generate scores for tumour microenvironment packages (e.g. ESTIMATE) All other analysis used R version 4.3.3. Samples were classified into the consensus molecular subtypes (CMS) using the CMS classifier package (1.0.0)^22^ by the random forest method and pathway derived subtypes (PDS) using the PDS classifier (v1.0.1)^2^. Single sample gene set enrichment analysis (ssGSEA) scores were generated using the GSVA package (1.50.5)^23^ (RRID:SCR_021058), the scores were generated using the gsva() function, with “ssgseaParam”and normalize = F. If subsequently scaled, this was performed using the scale function (default settings) in R with the gene sets as columns. Gene set enrichment analysis (GSEA) was carried out using fgsea (1.28.0)^24^ (RRID:SCR_020938) and enrichplot (1.22.0)^25^. The gene sets used were retrieved from the Molecular Signatures Database (MSigDB) package, msigdbr (7.5.1)^26^ (RRID:SCR_022870) as well as literature (Macrophage phenotypes)^8,27^. The two packages used to generate heatmaps were ComplexHeatmap (2.18.0)^28^ (RRID:SCR_017270) and circlize (0.4.16)^29^ (RRID:SCR_002141). The boxplots were created using ggplot2 (3.5.1)^30^ (RRID:SCR_014601) and statistically annotated by ggpubr (0.6.0)^31^ (RRID:SCR_021139). The statistical test used was Wilcoxon Rank Sum test with Hot PRS subgroup as the reference group. The microenvironment scores were generated using xCell (1.1.0) (rnaseq=F) and ESTIMATE (1.0.13). Intestinal stem cell index was calculated using ISCindex (0.0.0.9)^9^. Differential gene expression analysis was carried out on limma (v3.58.1)^32^ (RRID:SCR_010943) between CMS1 and CMS4 (adjusted p value < 0.05, n=6841 genes) and overlapped with the regenerative stem cell signature (n=232), resulting in a CMS1 RSC (n=61) and CMS4 RSC(n=51) gene signature. For KPN organoids feature counts were downloaded from https://www.ebi.ac.uk/biostudies/arrayexpress. KPNs were treated with either Interferon gamma (E-MTAB-11769) or TGF-β (E-MTAB-11784). These feature counts were then normalized using DESeq2. Within the design matrix, both genotype and condition were merged to form a ‘group’ column (e.g. KPNUT, KPTTGFB). Genes with a total read count below 10 across all samples were removed. VST transformation was then preformed with blind = F for both analyses, with the KPN samples selected for visualization.

### Transcriptional signature generation

The PRS signatures were generated using limma (3.58.1), the differential genes were ordered by logFC, with the top n=50 genes between either poly(I:C) treated epithelial or macrophage relative to their control counterparts (removing rows that included more than one gene name). The two signatures were found to have n=24 genes in common which created an Overlap PRS, with the remaining genes creating two unique signatures, epithelial PRS (n=26) and macrophage PRS (n=26). The PRS score for each signature was the mean expression value of genes in that specific signature. The PRS sum score, was the score for each signature added together.. Omnipath (3.10.1) was used to access the CollecTRI resource, which holds information on transcription factors, including the direction in which they regulate their target genes. This was used in combination with a univariate linear model (run_ulm) function from decoupleR (2.8.0) package to calculate transcription factor activity scores, with significant transcription factors found for epithelial and macrophage (relative to their control counterparts), having Wilcoxon rank sum test p value < 0.05. PHATE analysis (phateR v1.0.7)^33^ was utilised to visualise the biological trajectory based on the PRS genes in GSE39582 with Hot/Warm/Cold and CMS annotation on samples.

### Bioinformatic analysis, single cell data

Pre-processed scRNA-seq data (GSE132465; SMC cohort; n=23 CRC patients)^34^ were downloaded into a Seurat object. Following Seurat pipeline (RRID:SCR_016341), n=23,075 genes were kept after gene filtering (0 counts per <10 cells excluded) and SCTransform was applied for normalisation. To select for macrophage populations, within the myeloid populations from tumour samples, conventional dendritic cells (cDCs) and “Unknown” were removed, only keeping cells labelled as “Pro-inflammatory”, “Proliferating” and “SPP1+” in Seurat object (n=5586) for downstream single-cell analyses. Dimensionality reduction methods, including PCA and UMAP (dims=1:20), was performed using Seurat R package (5.0.1)^35^. To generate single cell-based enrichment scores on gene signatures, including interferon responses (Hallmarks), M1/M2 phenotypes and PRS, enrichit function from escape R package (1.6.0)^36^ was used and visualised on UMAP generated from M1/M2 genes with cells displaying top 25% enrichment scores as gradient using scCustomize R package (2.0.1)^37^ (RRID:SCR_024675). M1 and M2 macrophages were defined using the M1/M2 signature scores and divided into tertiles: High, Mid and Low. Spearman correlation was visualised as correlation heatmap using corrplot (0.92) (RRID:SCR_024683).

### Statistical analysis

Analysis carried out in the laboratory were compared using t-test within Prism (RRID:SCR_002798), otherwise direct comparisons were carried out using Wilcoxon rank sum test, using ggpubr (0.6.0) (RRID:SCR_021139). Pearsons correlation was carried out using cor.test() function within the stats package (4.3.3) (RRID:SCR_025968).

### Data availability

The publicly available bulk and single cell gene expression dataset used in this study for both the human and mouse models is referred to in the Methods, with corresponding GEO accession numbers. The new poly(I:C)-treated datasets from HCT116 and THP-1 lineages are available on GEO, under accession number GSE28054.

## Author Contributions

**SMC:** Conceptualization, Methodology, Formal analysis, Investigation, Data Curation, Writing original draft.

**SS:** Conceptualization, Methodology, Formal analysis, Investigation, Data Curation, Writing original draft.

**NEM:** Methodology, Formal analysis, Investigation, Data Curation.

**SBM:** Formal analysis, Investigation, Data Curation.

**RB:** Formal analysis, Investigation.

**AY:** Formal analysis, Investigation, Data Curation.

**RA:** Formal analysis, Investigation.

**CB:** Formal analysis, Investigation.

**AL:** Formal analysis, Investigation.

**KR:** Methodology, Formal analysis, Investigation, Data Curation.

**TL:** Formal analysis, Investigation, Data Curation.

**RR:** Resources, Supervision.

**FRT:** Writing, Supervision, Project administration, Data Curation.

**NCF:** Formal analysis, Investigation.

**TM:** Resources, Supervision.

**ML:** Resources, Supervision.

**AC:** Resources, Supervision.

**SJL:** Resources, Supervision.

**AER:** Resources, Supervision.

**DBL:** Resources, Supervision, Funding acquisition.

**DS:** Conceptualization, Methodology, Formal analysis, Investigation, Resources, Supervision, Data Curation.

**OJS:** Conceptualization, Resources, Supervision, Funding acquisition.

**PDD:** Conceptualization, Methodology, Formal analysis, Investigation, Resources, Supervision, Data Curation, Funding acquisition, Writing original draft.

## Acknowledgements and Financial Support

This work was supported by CRUK early detection grant (A29834), CRUK International accelerator programme, ACRCelerate, (A26825), MRC National Mouse Genetics Network programme (MC_PC_21042), US–Ireland R01 award (HSCNI, STL/5715/15)). General support for the Dunne research group via the QUB Foundation.

## Figure Legends

**Supplementary Figure 1:**
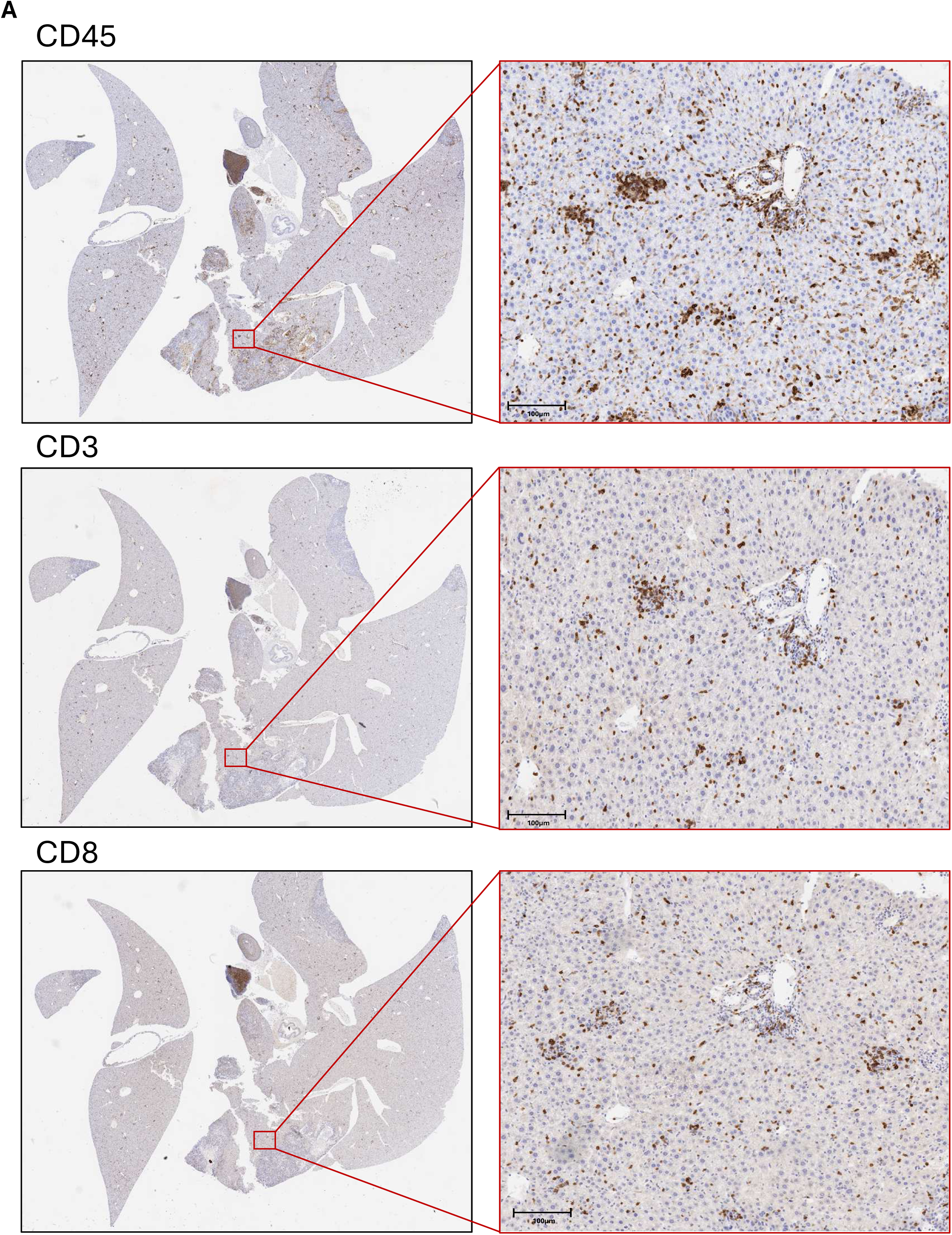
Immunohistochemistry staining of liver metastasis from KPN mice. **(A)** Immunohistochemistry staining of liver sections from KPN mice treated with poly(I:C) of CD45, CD3 and CD8.

**Supplementary Figure 2:**
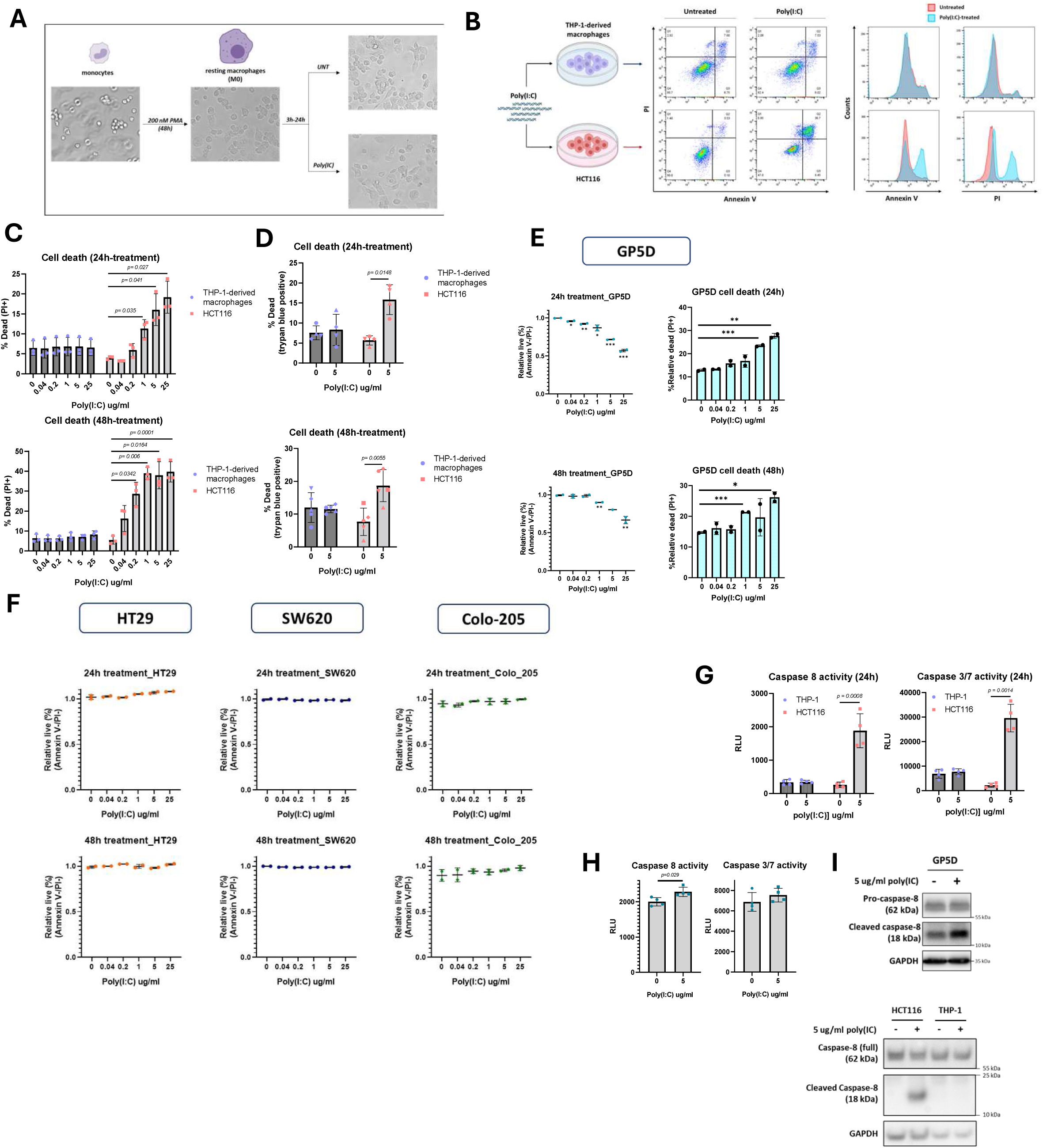
Cell lineage specific response to poly(I:C). **(A)** Differentiation process of monocytes into resting macrophages. **(B)** Cell death quantified using flow cytometry. **(C-D)** Cell death quantification across different poly(I:C) doses at 24 and 48hrs post poly(I:C) treatment (Student t-test, carried out on Prism). **(E)** Cell death quantification at 24hrs (top) and 48hrs (bottom) following the administration of poly(I:C) to GP5D cells. **(F)** Cell death quantification at 24hrs (top) and 48hrs (bottom) of HT29, SW620 and Colo-205 epithelial cell lines. **(G)** Caspase 8 and Caspase3/7 activity at 24hrs in treated and untreated macrophage (THP-1) and epithelial (HCT116). **(H)** Caspase 8 and Caspase 3/7 activity in poly(I:C) treated and untreated GP5D cells. **(I)** Pro- and cleaved-caspase 8 in poly(I:C) treated and untreated GP5D cells (top) and poly(I:C) treated and untreated macrophage (THP-1) and epithelial (HCT116) (bottom). (Student t-test carried out on Prism: ns: p > 0.05, *: p <= 0.05, **: p <= 0.01, ***: p <= 0.001, ****: p <= 0.0001). Error bars = 1 standard deviation.

**Supplementary Figure 3:**
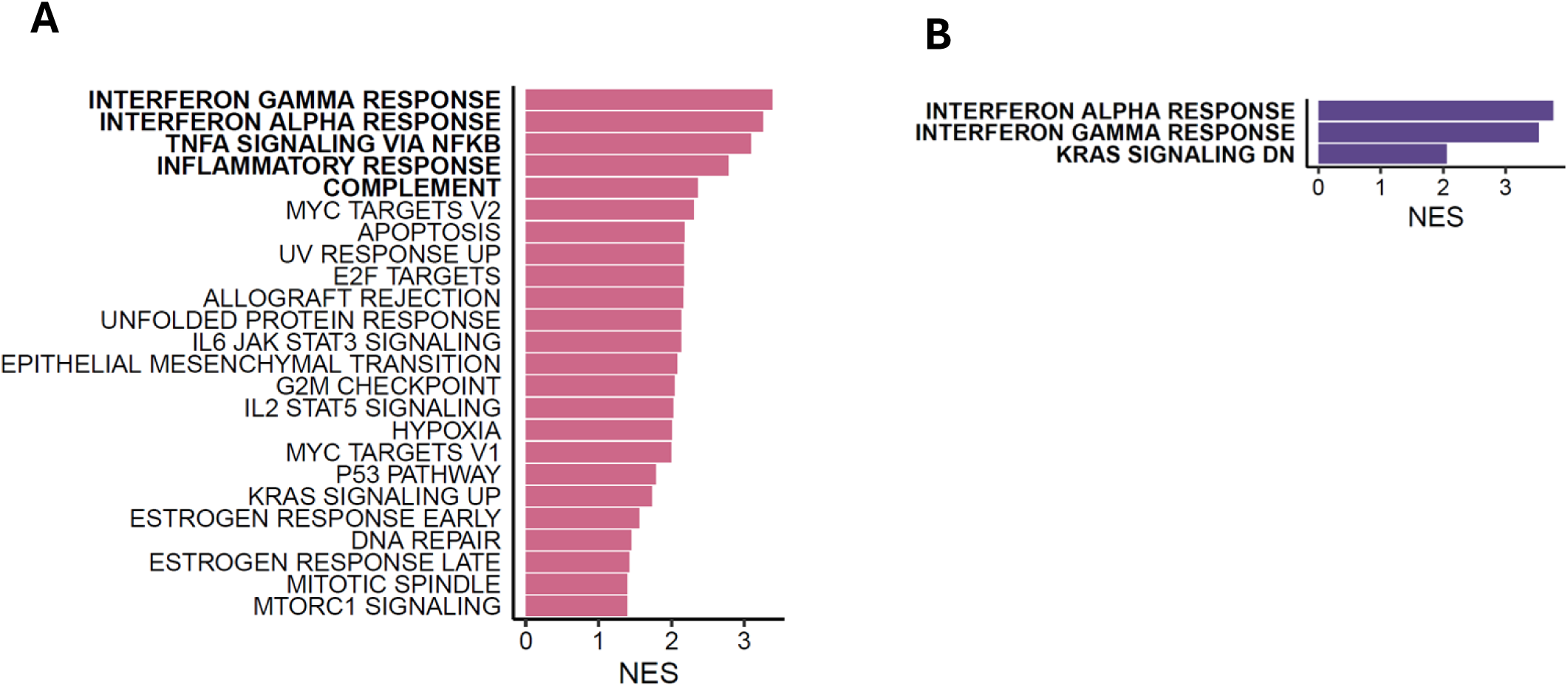
Transcriptional analysis of *in vitro* model. **(A)** Gene set enrichment analysis (GSEA) of the significant Hallmark gene sets (adjusted p value < 0.05) in epithelial and **(B)** macrophage cells treated with poly(I:C) relative to their untreated controls, organised by normalized enrichment score with the top n=5 (n=3 for macrophage due to low number of significant gene sets) in bold font.

**Supplementary Figure 4:**
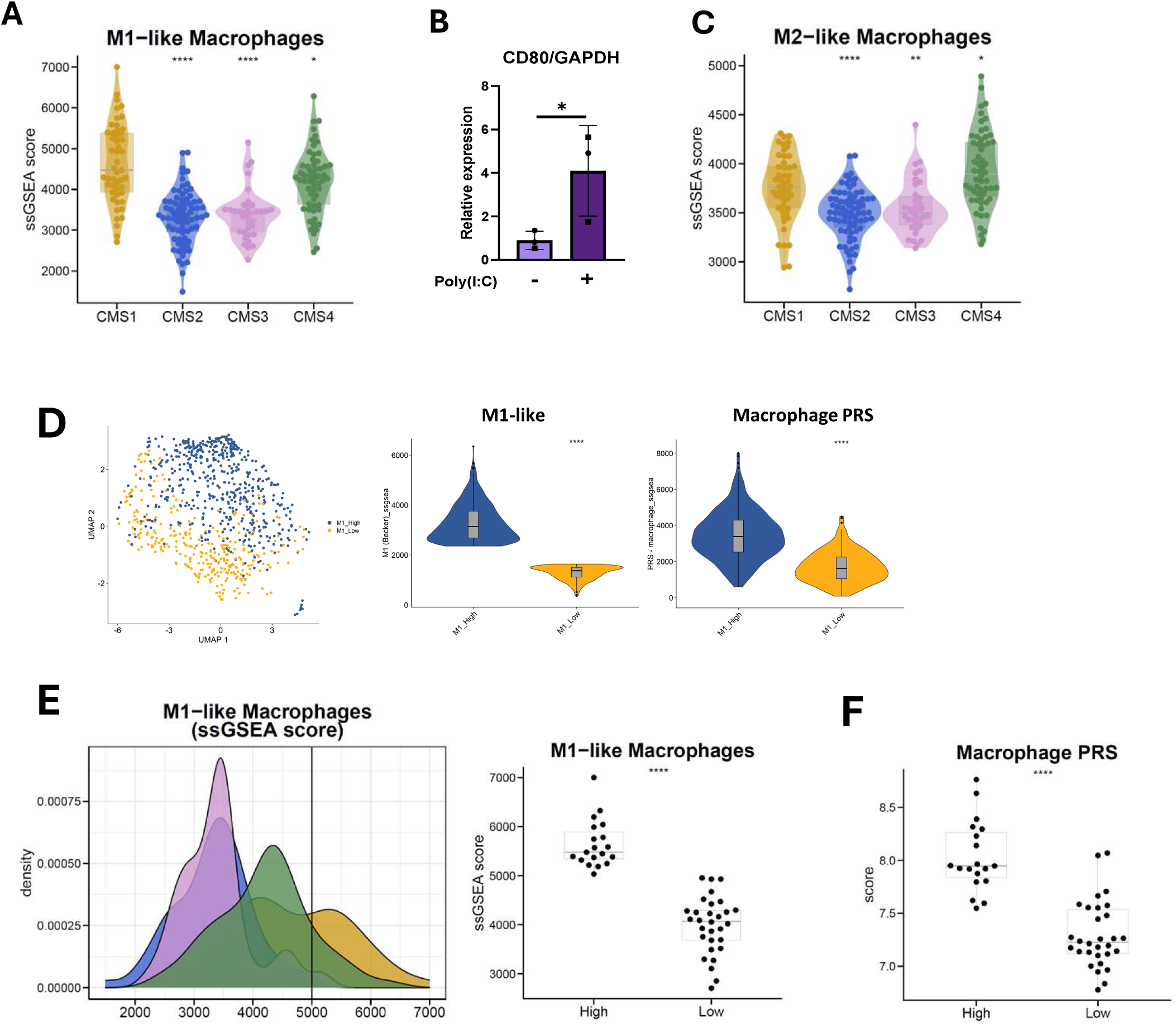
Poly(I:C) treatment induces an M1-like phenotype that is encapsulated in the macrophage PRS. **(A)** M1-like transcriptional signature single sample gene set enrichment analysis (ssGSEA) scores across the CMS’s (GSE39582) (n=258) (Wilcoxon rank sum test, CMS1 as reference group). **(B)** CD80 quantitative polymerase chain reaction (qPCR) between the poly(I:C) treated and untreated macrophages (Student t-test carried out on Prism: p <= 0.05). Error bars = 1 standard deviation. **(C)** M2-like transcriptional signature single sample gene set enrichment analysis (ssGSEA) scores across the CMS’s (Wilcoxon rank sum test, CMS1 as reference group). **(D)** The myeloid subpopulation from the single cell RNA-sequencing cohort (GSE132465) (n=23) that were split into tertiles based on M1 and M2 transcriptional scores, with the top (M1 high) and bottom (M1 low) tertile visualised (left) and compared based on their M1-like transcriptional score and macrophage PRS score (right) (Wilcoxon rank sum test). **(E)** M1-like transcriptional signature ssGSEA scores across the CMS’s visualised as a ridge plot (left), with a vertical line creating a stratification point into M1 high and M1 low of CMS1 samples (right). **(F)** The macrophage PRS in the M1 high subset of CMS1 samples compared to the M1 low subset of CMS1 samples. (Wilcoxon rank sum carried out by ggpubr: ns: p > 0.05, *: p <= 0.05, **: p <= 0.01, ***: p <= 0.001, ****: p <= 0.0001).

**Supplementary Figure 5:**
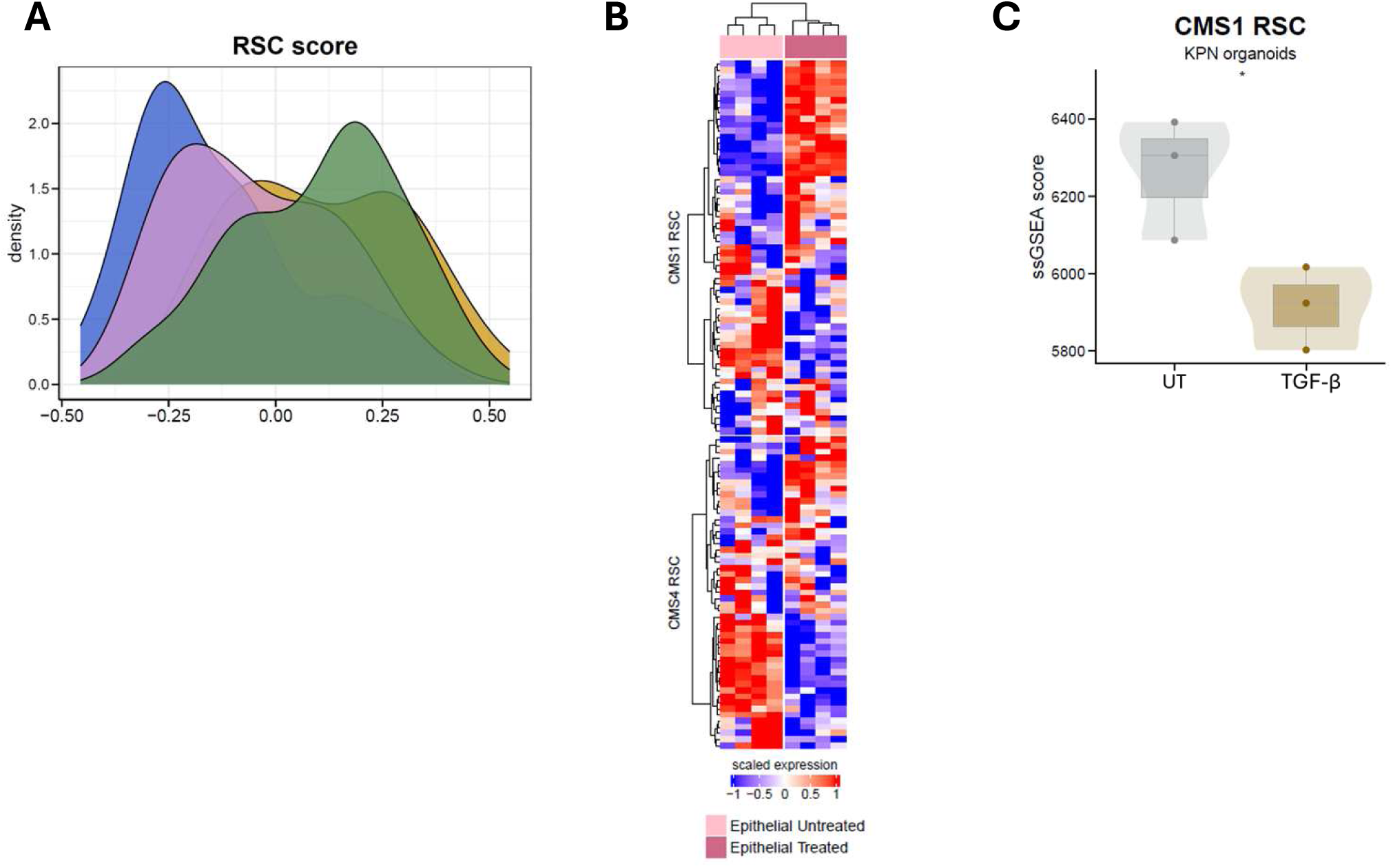
Poly(I:C) induces a CMS1-like regenerative stem phenotype. **(A)** Regenerative stem cell (RSC) signature score across the CMS’s visualised in a ridge plot (GSE39582) (n=258). **(B)** The genes from the CMS1 RSC and CMS4 RSC visualised across the poly(I:C)-treated and untreated epithelial cells. **(C)** ssGSEA scores of CMS1 RSC in KPN Organoids treated with TGF-β (E-MTAB-11784) (t-test).

**Supplementary Figure 6:**
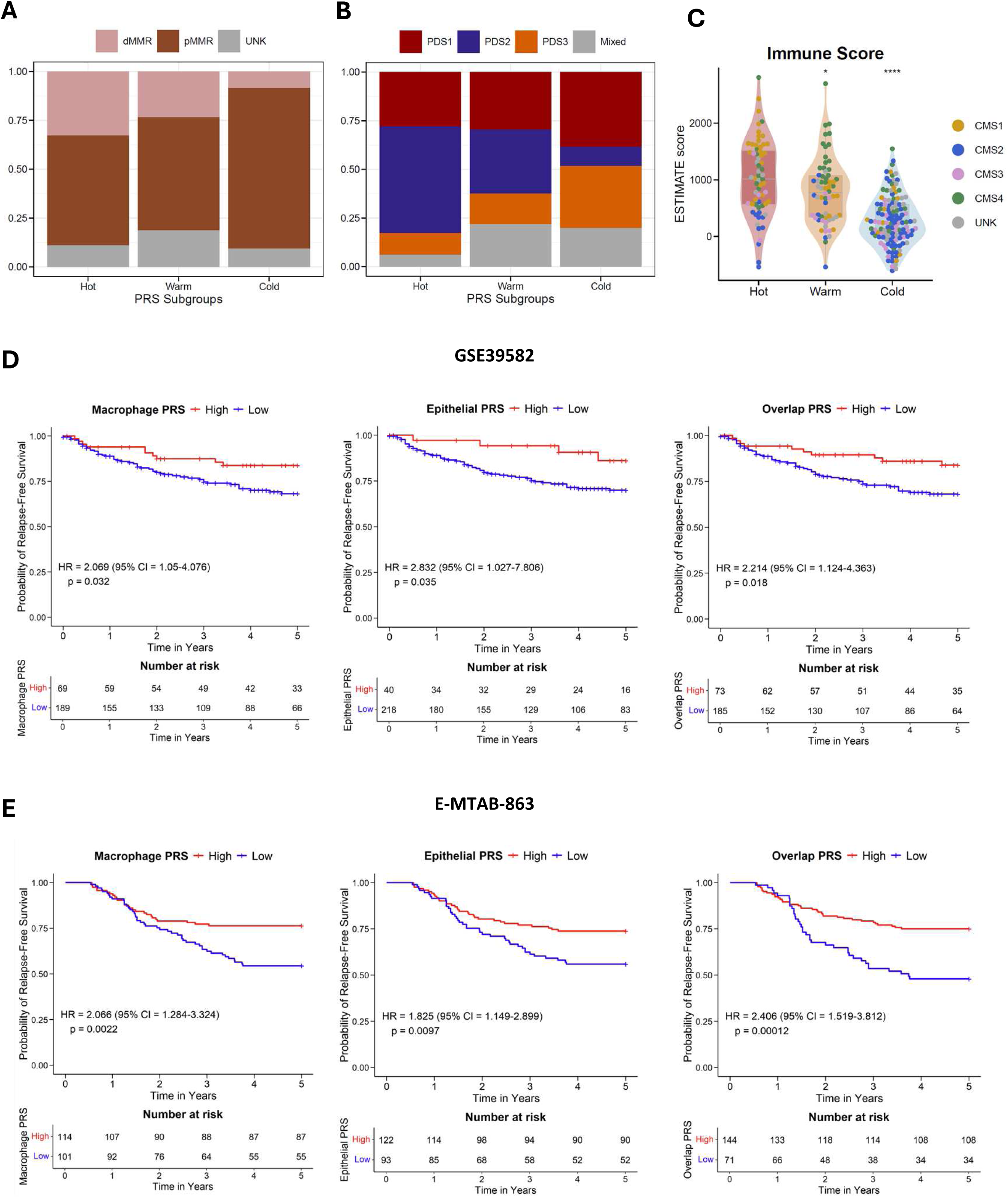
Poly(I:C) biology aligns with an immune active CMS1-like phenotype. **(A)** Proportion of mismatch repair deficient (d) and proficient (p) samples across the PRS subgroups (GSE39582) (n=258). **(B)** Proportions of samples classified as the pathway derived subtypes (PDS) within the PRS subgroups. **(C)** ESTIMATE immune score across the PRS subgroups (Wilcoxon rank sum test with Hot subgroup as reference). **(D)** Kaplan-Meier survival estimates of macrophage, epithelial and overlap PRS in the GSE39582 cohort (n=258) and **(E)** an independent stage II colon cancer cohort (n=215) (E-MTAB-863) (all significant, p value < 0.05). (Wilcoxon rank sum carried out by ggpubr: ns: p > 0.05, *: p <= 0.05, **: p <= 0.01, ***: p <= 0.001, ****: p <= 0.0001).

